# Optimization of DNA extraction from human urinary samples for mycobiome community profiling

**DOI:** 10.1101/503946

**Authors:** A. Lenore Ackerman, Jennifer T. Anger, Muhammad Umair Khalique, James E. Ackerman, Jie Tang, Jayoung Kim, David M. Underhill, Michael R. Freeman, NIH Multidisciplinary Approach to the Study of Chronic Pelvic Pain (MAPP)

## Abstract

**Introduction:** Recent data suggest the urinary tract hosts a microbial community of varying composition, even in the absence of infection. Culture-independent methodologies, such as next-generation sequencing of conserved ribosomal DNA sequences, provide an expansive look at these communities, identifying both common commensals and fastidious organisms. A fundamental challenge has been the isolation of DNA representative of the entire resident microbial community, including fungi.

**Materials and Methods:** We evaluated multiple modifications of commonly-used DNA extraction procedures using standardized male and female urine samples, comparing resulting overall, fungal and bacterial DNA yields by quantitative PCR. After identifying protocol modifications that increased DNA yields (lyticase/lysozyme digestion, bead beating, boil/freeze cycles, proteinase K treatment, and carrier DNA use), all modifications were combined for systematic confirmation of optimal protocol conditions. This optimized protocol was tested against commercially available methodologies to compare overall and microbial DNA yields, community representation and diversity by next-generation sequencing (NGS).

**Results:** Overall and fungal-specific DNA yields from standardized urine samples demonstrated that microbial abundances differed significantly among the eight methods used. Methodologies that included multiple disruption steps, including enzymatic, mechanical, and thermal disruption and proteinase digestion, particularly in combination with small volume processing and pooling steps, provided more comprehensive representation of the range of bacterial and fungal species. Concentration of larger volume urine specimens at low speed centrifugation proved highly effective, increasing resulting DNA levels and providing greater microbial representation and diversity.

**Conclusions:** Alterations in the methodology of urine storage, preparation, and DNA processing improve microbial community profiling using culture-independent sequencing methods. Our optimized protocol for DNA extraction from urine samples provided improved fungal community representation. Use of this technique resulted in equivalent representation of the bacterial populations as well, making this a useful technique for the concurrent evaluation of bacterial and fungal populations by NGS.

## Introduction

Multiple organs, such as the gut, oral cavity, and vagina, have long been known to harbor communities of microbes that can protect against or contribute to disease under different circumstances. The urinary tract, however, was widely thought to be sterile until only recently, when extended culture techniques and the detection of microbial DNA definitively demonstrated microbial communities of great diversity within this site.[1-3] Currently, culture-independent microbial characterization using the sequencing of highly conserved DNA regions, such as the ribosomal RNA gene locus (rDNA), is widely-accepted as a useful, sensitive tool to explore microbial populations. These next-generation sequencing (NGS) technologies are particularly useful in characterizing microbes that may be difficult to culture or that are present in low abundance (the “rare biosphere”).[4] Therefore, the composition and diversity of the urinary microbiome has likely been drastically understated, in part, due to dependence on culture methods to identify resident species.

With the development of affordable, rapid, and scalable culture-independent methods for the study of bacterial communities, the last decade has seen a massive expansion in studies aimed at profiling commensal communities in a multitude of organs not included in the large-scale Human Microbiome Project (HMP), such as the urinary tract. Using NGS methods, multiple studies have demonstrated that perturbations in the urinary microbiota appear to correlate with Lower Urinary Tract Symptoms (LUTS).[5-13] The clinical significance and utility of these alterations, however, remain unclear, primarily due to challenges that persist for the characterization of microbes from low biomass specimens, such as urine.

Due to these limitations, we still lack vital information about the content of normal urine and its relationship to dysbiosis and/or disease. Studies examining the urinary microbiome thus far demonstrate wide variation in their ability to consistently detect microbial species. In many studies, approximately half of patient samples do not have bacterial sequences of sufficient quality for analysis[2, 6, 14]; in other studies, this efficiency could be improved with the use of multiple amplification steps[11], but this may introduce new biases that could skew results. This low sequencing efficiency is likely due to the combination of low biomass and the unique qualities of urine, which include high variability in osmolality/salt content, high abundance of PCR inhibitors, and fluctuating levels of cellular material, all in all making urine a challenging biological fluid to study. The question remains as to whether these sequence-negative samples are truly negative for microbes or whether our detection methods are inadequate to fully characterize these specimens. Until this question can be answered, it remains a very real possibility that the subset of samples analyzed, the “sequence-positive” group, may represent a unique subgroup within the analyzed population with higher microbial loads, whose findings cannot be generalized to the larger sample population.

Even less is known about the composition of non-bacterial populations, such as fungi, viruses, archaea, and protozoa, in the genitourinary tract and other human organs, primarily from a lack of well-researched tools for their analysis. Despite these challenges, alterations in the fungal microbiota (the “mycobiome”) in the absence of frank infection have been demonstrated in multiple human diseases, such as hepatitis [15], atopic dermatitis [16], inflammatory bowel disease [17-19], cystic fibrosis [20], allergy/atopy [21], asthma [22], and psoriasis [23, 24]. As yet, only a few analyses have examined aspects of the urinary mycobiome. *Candida spp*. have been detectable in urinary samples by culture,[5-8] demonstrating their viability. Fungi were also detectable in urine from patients with urological chronic pelvic pain syndromes (UCPPS) using the targeted Ibis T-5000 Universal Biosensor system.[25] Interestingly, fungi were detected more frequently in UCPPS patients during symptomatic flares, while no significant differences in the bacterial microbiota could be identified, implicating fungi as important players in lower urinary tract symptomatology. Even in this culture-independent study, however, fungi were detected in less than 10% of patients overall. Again, it is unclear if this low number is representative of the absence of fungi in the majority of subjects or represents severe limitations in our current technologies.

Further progress in identifying consistent microbial markers or understanding the pathophysiology of microbial interactions in the urinary tract requires methodologies that adequately and reliably characterize these populations, and which include fungi and other microbes in addition to bacteria. In this study, we sought to identify the most effective strategies for extracting and identifying microbial DNA from urine, with a focus on enhancing the detection of fungi. Using an iterative approach, we optimized urine sample processing at multiple steps to increase DNA yields and population representation to generate more consistent data from sequencing-based microbial population analyses.

## Materials and Methods

This study was approved by the Cedars-Sinai Institutional Review Board (Pro00033267) and written consent was obtained from all subjects.

### DNA yield assessment

Overall DNA yields and quality (assessed by OD_260_/OD_280_ ratios) were measured on the NanoDrop 2000 Spectrophotometer (Thermo Scientific). Fungal DNA levels were assessed in duplicate by quantitative Real-Time Polymerase Chain Reaction (qRT-PCR) analyses on a Mastercycler Realplex2 (Eppendorf) using the SYBR Green PCR kit as instructed by the manufacturer (Applied Biosystems). Fungal levels were assessed using the Fungiquant primers (forward: 5’-GGRAAACTCACCAGGTCCAG-3’; reverse: 5’- GSWCTATCCCCAKCACGA-3’)[26] that recognize a highly-conserved segment of the fungal 18S rDNA region, while bacterial levels were assessed using 16S rDNA primers (forward: 5’-ACTCCTACGGGAGGCAGCAGT-3’; reverse: 5’-ATTACCGCGGCTGCTGGC-3’), a universal primer with broad specificity for bacteria. The qRT-PCR protocol employed an initial denaturation at 94°C for 10 min, followed by 35 cycles of denaturation at 94°C for 30 s, annealing at 55°C for 30 s, and elongation at 72°C for 2 min, followed by an elongation step at 72°C for 30 min. Relative quantity of bacterial and fungal DNA yields, consistent from experiment to experiment, was calculated by the comparative CT method (2^−ΔΔC^T method)[27] and normalized to a DNA standard curve derived from a mixed bacterial and fungal culture that remained constant over all tests. Samples with greater than 3% variance between duplicates were reanalyzed in duplicate. An aliquot of 1 μl of the PCR product was evaluated by 2% agarose gel electrophoresis.

### Evaluation of Individual protocol enhancements

For the initial, iterative analyses, specimens were obtained from mid-stream urine collections from multiple male and female subjects, all of whom denied any urinary symptoms, after preparation of the external urethral meatus with chlorhexidine gluconate wipes. Urine specimens were mixed well, then divided into 1 ml samples and centrifuged at 5000 relative centrifugal force (rcf) to pellet cellular material prior to parallel processing to test the individual protocol variations described below.

#### Enzymatic disruption

Sample pellets were initially resuspended in 500 μl enzyme buffer (0.5 M Tris, 1mM EDTA, and 0.2% 2-mercaptoethanol, pH 7.5). We added 200 U/ml Lyticase (Sigma Aldrich), 20 mg/ml Lysozyme (Thermo Scientific), both enzymes, or buffer alone without enzyme. Samples were then incubated for 30 min. at 30°C, with inversion of the tubes every 5-10 min. Subsequently, samples were centrifuged at 1500 rcf for 5 min, the supernatant removed, and the pellet resuspended in 800 μl of Stool DNA Stabilizer (Stratec Biomedical).

#### Mechanical disruption

Physical disruption of cell walls was accomplished with bead beating. The 800 μl post-enzymatic digestion cell suspension was transferred to a 2 ml centrifuge tube containing 100 μl 0.1 mm and 300 ul 0.5 mm silica beads (Biospec Products, Inc.). Samples were agitated twice for 1 min each on a standard Vortex mixer using a Vortex Adapter for bead beating (MO BIO Laboratories Inc.). Samples were centrifuged for 15 s at 17000 rcf between bead beating periods.

#### Thermal disruption

Samples were heated to 95°C for 10 min, with a brief vortex to ensure adequate mixing 5 min into the incubation. After a second, brief vortexing step, samples were incubated on ice (0°C) for 5 min, then centrifuged for one min. at 17000 rcf after each boil/freeze cycle.

#### Proteinase digestion

After cell wall disruption, cell lysates were transferred to new tubes containing an equal volume of buffer AL (Qiagen) containing varying concentrations of Proteinase K (0, 12, 24, 48, 72, 96, 120, or 144 mAU/ml)(Qiagen), then incubated at 70°C for 10 min.

#### Addition of carrier DNA

When specified, polyadenylic acid carrier DNA (PolyA) (Roche Diagnostics) was added to the cell lysates at the time of proteinase K digestion.

#### Column DNA extraction

Following the specified disruption and digestion steps, 250 μl 100% Ethanol was added and briefly mixed by vortexing, prior to applying the cell lysates to Qiagen Mini DNA Spin columns (Qiagen). The columns were washed twice with a column volume of buffer AW (Qiagen) by centrifugation at 17000 rcf for 1 min and residual alcohol removed with a third spin without wash buffer. DNA was then eluted from the column in 60 μl warm Tris-EDTA (TE) buffer.

### Confirmation of protocol components in aggregate analysis

For 4 subjects (2 male and 2 female), >60 ml of urine were obtained and divided into 1 ml samples within 1 h of sample collection. Twenty unique conditions were analyzed following centrifugation at one of three centrifugation conditions: 1) 1500 rcf for 15 min., 2) 5000 rcf for 20 min., or 3) 16000 rcf for 10 min. The resulting pellets were frozen and stored at −80°C. 20 unique combinations of the conditions explored in initial, iterative analysis were performed, with inclusion or exclusion of the individual enzymatic treatments, mechanical and thermal disruption steps, and proteinase digestion in almost all combinations. For this panel of conditions, carrier DNA was included in all samples to provide better discrimination of differences in these low volume samples.

Relative DNA yields for each condition were determined by fungal-specific qPCR as specified above; for each sample, yields were scaled to equal variance for all samples to allow plotting of the median yields for each condition as a heat map.

### Determination of optimal sample volume

Large volume urine samples (>100 ml) from 3 male and 3 female subjects were mixed well and subdivided into 1, 2.5, 5, 10, 25, and 50 ml aliquots. Each sample was centrifuged at 1500 rcf and the supernatant decanted. After pelleting, all samples were identically processed using the optimized protocol detailed above. All aliquots for an individual subject were processed in batches to minimize batch-to-batch variation. The resulting fungal DNA concentrations were then quantitated by qRT-PCR. Taxal diversity was also examined by 2% agarose gel electrophoresis.

### Sample subdivision and pooling

To evaluate if processing lysates in smaller volumes provided increased DNA yields, samples were subdivided into smaller aliquots after mechanical disruption. Seven identical urine specimens were pelleted, digested with lysozyme and lyticase, then subjected to bead beating with a mixture of silica beads as detailed above. Sample quantities ranging from 100 μl to 400 μl (of an approximately 500 μl total lysate volume) at 50 μl intervals were aspirated off of the beads and subjected to thermal disruption, proteinase K digestion and DNA-column binding and elution.

To examine if total DNA yields could be increased by pooling these smaller aliquots, sample lysates were subdivided into two 250 μl aliquots after mechanical disruption, then subjected to thermal disruption and proteinase digestion separately. These two samples were then applied to either a single DNA-binding column in succession or to two separate columns, eluted and pooled after elution. Overall and fungal-specific DNA yields were then measured using NanoDrop DNA quantitation and fungiquant qRT-PCR.

### Light microscopy

After centrifugation, cellular pellets from urine were resuspended in 5 ml PBS and mixed well with a pipette. A 10 μl aliquot was transferred to a 75 × 26-mm glass slide and covered with an 18 × 18-mm coverslip, ensuring that the sediment was uniformly distributed but not escaping from the edges of the coverslip. Using an inverted IX51 microscope (Olympus), images without staining were captured at ×400 (objective lens 40× in combination with wide field 10× eyepiece) to generate a field area of 0.196 mm^2^.

### Comparison with commercial methods

We compared our optimized approach to three, commonly used commercial kits for DNA extraction: PSP^®^ Spin Stool DNA Plus Kit (Stratec Biomedical), PureLink™ Microbiome DNA Purification Kit (ThermoFisher Scientific), and QIAamp DNA Stool Mini Kit (Qiagen). Large volume urine specimens (>120 ml) from 9 subjects were divided into four 30 ml specimens and pelleted by centrifugation at 1500 rcf. Mid-vaginal swabs were obtained from female subjects using FloQSwabs (Copan Diagnostics). Swabs were gently agitated for 30 min in 500 μl enzyme buffer (0.5 M Tris, 1mM EDTA, and 0.2% 2-mercaptoethanol, pH 7.5), before removing the swab; the resulting cell suspension was then processed as for urine specimens. The identical urine samples and vaginal swabs were processed according to the manufacturers’ protocols for each kit or using our optimized protocol. Fungal and bacterial DNA yields in the eluents were then assessed by qPCR as specified above.

### Microbial Sequencing Analysis

#### Library Generation

DNA was isolated from urine using the specified protocols as described above. Fungal ITS1 and bacterial 16S regions amplicons were generated by PCR using the primers below modified to include Nextera XT v2 barcoded primers (Illumina) to uniquely index each sample. PCR reactions utilized Platinum SuperFi DNA Polymerase (Invitrogen) according to the following protocol: initial denaturation at 94°C for 10 min, followed by 40 cycles of denaturation at 94°C for 30 s, annealing at 48°C for 30 s, and elongation at 72°C for 2 min., followed by an elongation step at 72°C for 30 min.

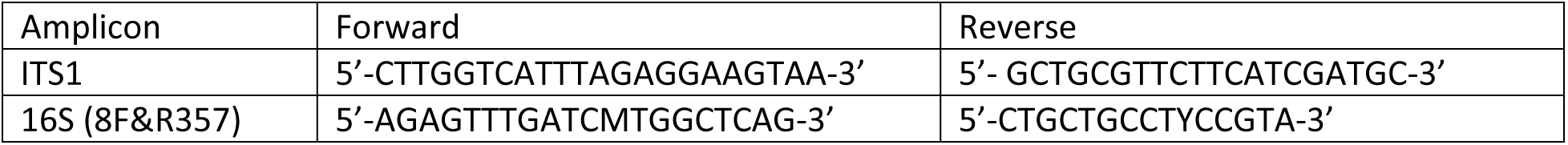

#### Next-generation sequencing

Amplicons generated above were sequenced at 2×300 paired-end sequencing on the Illumina MiSeq sequencer, according to manufacturer’s instructions. Raw data processing and de-multiplexing was performed using on-instrument MiSeq Reporter Software v2.6 as per manufacture recommendations. Demultiplexed 16S sequence data were processed and analyzed as previously described including OTU assignment by alignment to the GreenGenes reference database (May 2013 release) at 97% identity.[28] For analysis of ITS1 sequence data, raw FASTQ data were filtered to enrich for high quality reads including removing the adapter sequence by Cutadapt v1.4.1,[29] truncating reads with average quality scores less than 20 over a 3-base pair sliding window and removing reads that do not contain the proximal primer sequence or that contain a single unknown base. Filtered pair-end reads were then merged with overlap into single reads using SeqPrep v1.1 wrapped by QIIME v1.9.1.[30] Processed high-quality reads were then aligned to previously observed host sequences (including rRNA and uncharacterized genes in human) to deplete potential contamination. Operational taxonomic units (OTU) were identified by alignment of filtered reads to the Targeted Host Fungi (THF) custom fungal ITS database (version 1.6), [31] using BLAST v2.2.22 in the QIIME v1.9.1 wrapper with an identity percentage ≥97%.

### Diversity analysis

We performed rarefaction analysis. The original OTU table was randomly subsampled (rarefied) to create a series of subsampled OTU tables. Alpha diversity was calculated on each sample using the OTU table and a variety of metrics (chao1, observed species, etc.). The results of the alpha diversity were collated into a single file and the number of species identified for each sample versus the depth of subsampling was plotted. Shannon diversity indices were selected to show composite readout of microbial population evenness and richness.

### Statistical analysis

Differences in DNA yields between groups were compared using a two-tailed, paired Student’s *t* test with a 95% confidence interval. Data are presented as means ± SEM, unless otherwise stated. Statistical analyses were performed using Microsoft Excel 2016 (version 1803) or RStudio version 3 as appropriate.

## Results

### Sequential optimization of fungal DNA extraction

To optimize the procedure of isolating urinary microbial DNA, we began with a protocol described in the initial isolation of bacteria from urine specimens,[5, 32]. Small volumes of urine were initially centrifuged to concentrate cells and microorganisms, then subjected to DNA extraction using a standardized kit involving DNA binding and elution from an affinity column.

To concentrate the rare cellular material present in urine, samples were centrifuged under three conditions previously described for the isolation of fungi from low biomass fluids.[32-34] Samples prepared with an initial centrifugation speed of 1500 relative centrifugal force (rcf) for 20 min. yielded fungal DNA levels at least 1.5-fold higher than those prepared at 5000 rcf for 10 min., while yields from those centrifuged at 16000 rcf for 10 min. were substantially lower (Figure 1A).

**Figure 1.**
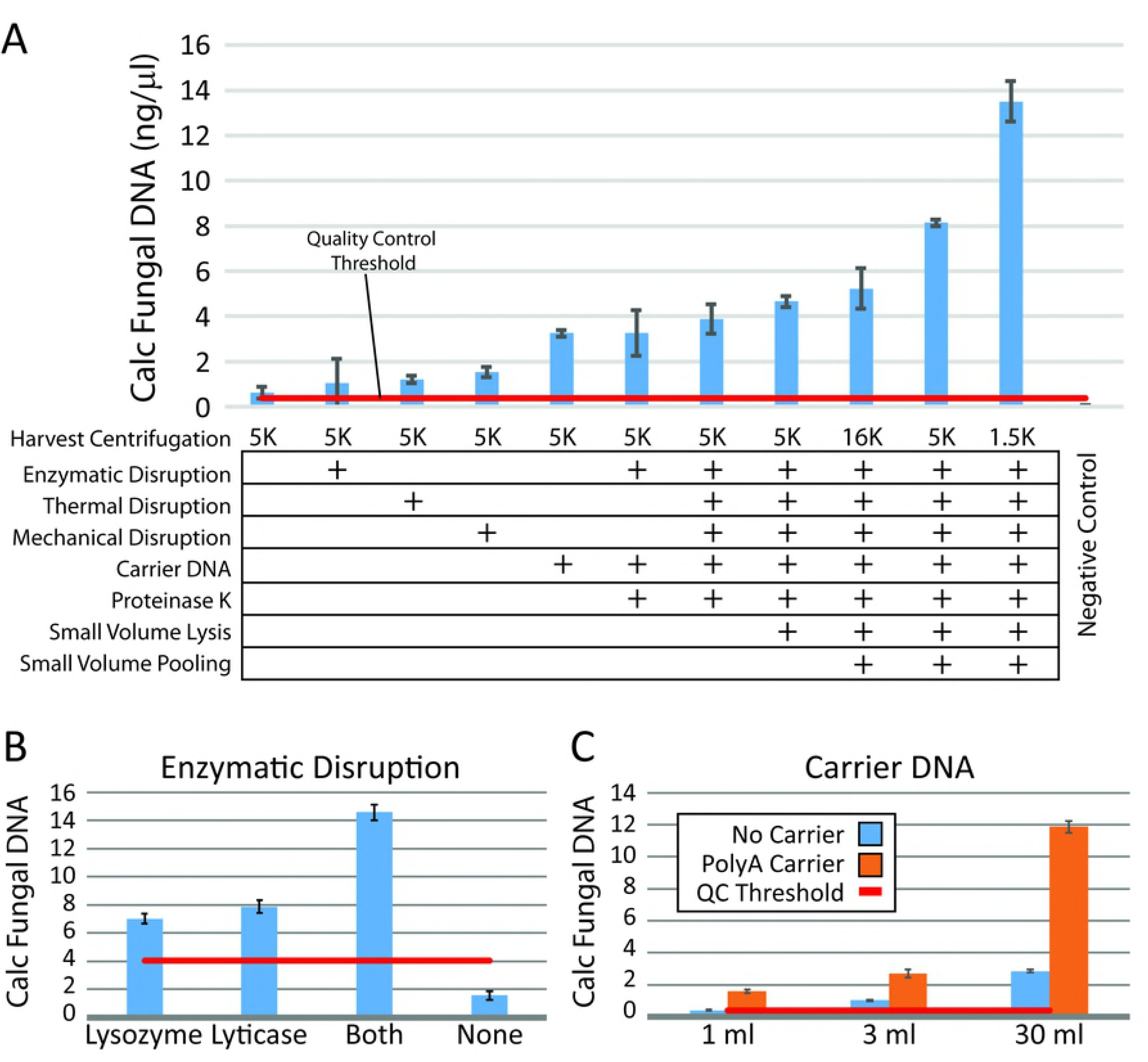
Optimization of Microbial DNA Extraction Requires Multiple Disruption Steps. (A) Eight variations in the protocol (at left) were noted to increase yields as determined by quantitative PCR. Relative fungal DNA yields were calculated from quantitative PCR using the Fungiquant pan-fungal PCR primer pair and normalized to a mixed fungal DNA standard. The negative control samples were processed in parallel, but did not have any input cellular material. Multiple protocol variations, such as enzymatic pre-digestion (B) or carrier DNA use during DNA column binding (C), were tested individually in triplicate for multiple subjects (minimum n=4), both male and female, before incorporating.

Fungi and some bacteria have cell walls, which can be resistant to digestion, leading to their absence or underrepresentation in culture-independent analyses. To optimize the isolation of organisms with robust cell walls, we examined the utility of an initial enzymatic digestion step to aid in cell wall dissolution.[35] Lysozyme, a glycolytic hydrolase that catalyzes the breakdown of peptidoglycan in gram-positive bacterial cell walls, is known to enhance gram-positive bacterial detection.[36] Lyticase, which hydrolyzes the poly- β(1➔3)-glucose present in yeast cell wall glycans, has been widely used in yeast DNA extraction, including PCR-based clinical assays.[34, 37] These enzymes were tested alone and in combination in comparison to omission of this step. Consistently, the combination of the two enzymes resulted in improved yields of both total DNA (data not shown) and relative fungal DNA levels calculated by qPCR (Figure 1B).

Particularly for fungi, physical disruption techniques, such as the thermal and mechanical steps described above, significantly improve fungal DNA purification,[38-40] again by further breaking down tough cell walls. Bead beating, which we performed using multiple sizes of silica beads, can be particularly useful in isolation of fungi such as Aspergillus, which is known to play a role in multiple human diseases.[41, 42] An additional thermal disruption step, with two freeze-boil (0°C/95°C) cycles, was also evaluated. Both methods used in isolation enhanced DNA extraction efficiency 2-3-fold over baseline (Figure 1A).

These disruption steps were followed by an additional digestion step with Proteinase K, a broad-spectrum serine protease, to remove any protein contamination and inactivate any remaining DNAase activity prior to cell and nuclear lysis. We tested a range of proteinase concentrations; while inclusion of the enzyme was important in enhancing DNA extraction efficiency, varying the proteinase concentration had much less effect. While a concentration of 24 mAU/ml (0.8 μg/ml) tended to provide the best results, a range of concentrations from 12-144 mAU/ml (4.8 μg/ml) did not differ significantly in their enhancement of DNA recovery (data not shown).

To maximize DNA recovery, we also evaluated the addition of carrier DNA. Because naturally occurring carriers, such as salmon sperm DNA, contain rDNA sequences with partial homology to other eukaryotic DNA, we chose a synthetic carrier, polyadenylic acid, which has shown efficacy in enhancing recovery of low abundance DNA from human biological samples.[43] Supplementation of carrier DNA increased both overall (data not shown)and fungal-specific DNA yields 2-4 fold across all samples (Figure 1C). The best combination of all techniques tested resulted in an almost 14-fold increase in fungal DNA yields, comprising an optimal protocol utilizing low-speed centrifugation, enzymatic, mechanical, and thermal cell wall disruption, inclusion of carrier DNA, and proteinase K digestion in combination.

### Confirmation of protocol on standardized samples

Each of the individual conditions noted to increase DNA yields were tested in aggregate on a panel of urine specimens from both male and female patients. Large volume (>75 ml) urine specimens from 4 subjects (2 male and 2 female) were divided into small equal aliquots (1 ml), and then processed in parallel to confirm the enhancement of DNA purification with the modifications observed in the individual experiments detailed above. This larger-scale optimization panel assessed the variations in cell wall disruption methods (thermal and mechanical), enzymatic pre-treatment methods (lysozyme and lyticase), proteinase K digestion, and centrifugation speed in almost all combinations (Figure 2). Calculation of the relative fungal DNA yields from these 60 variations in isolation methodology revealed a clear pattern, with improved yields resulting from the optimized protocol defined above with multiple disruption methods, combined enzymatic digestion, and lower centrifugation speeds.

**Figure 2.**
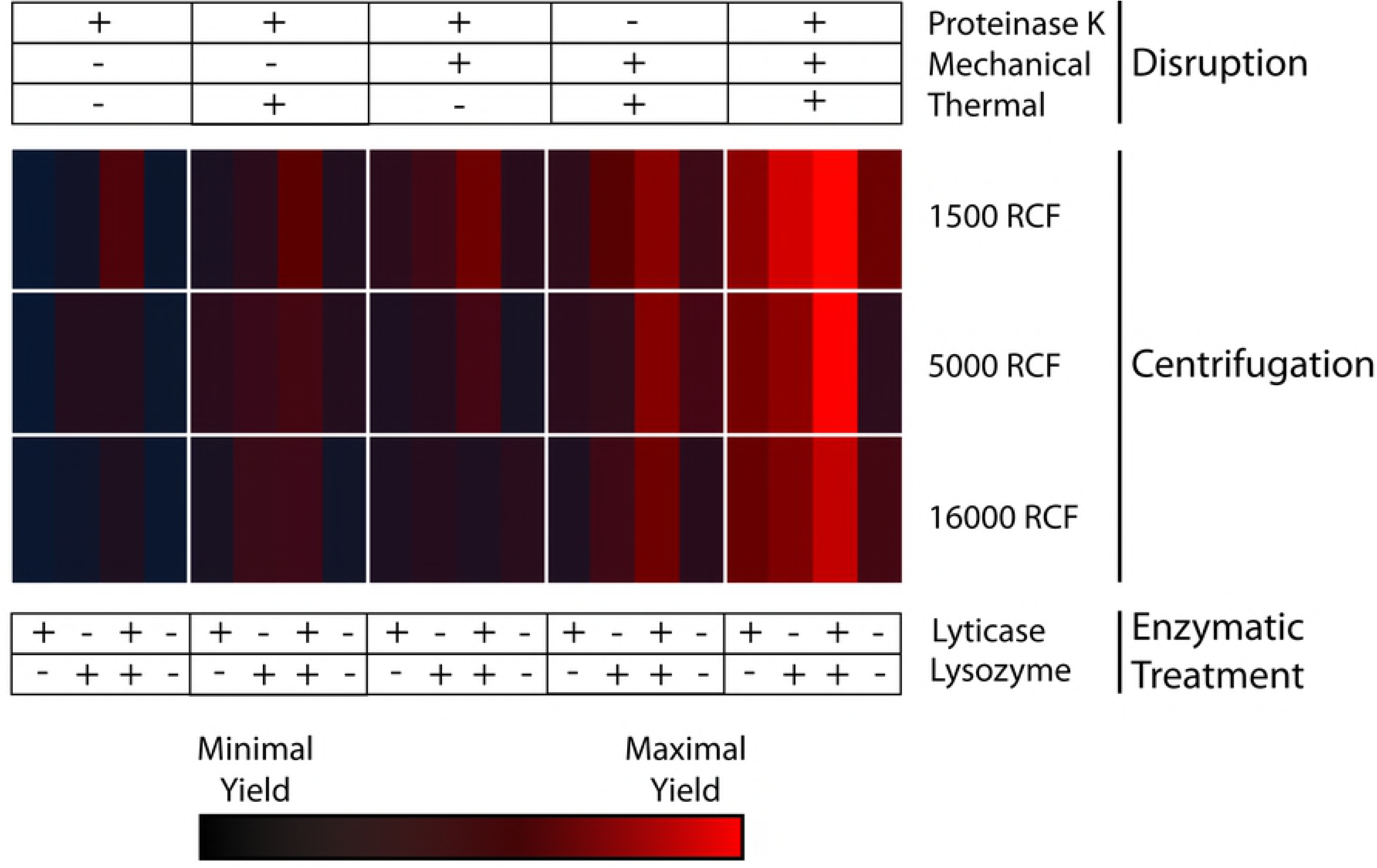
Large-Scale Confirmation of Optimization of Microbial DNA purification. The individual conditions noted to increase yields were tested in aggregate in a larger-scale optimization panel. DNA was concurrently isolated from 60 identical 1 ml urine samples from each of 4 subjects with variations in cell wall disruption methods (as indicated at the top), enzymatic pre-treatment methods (bottom), and centrifugation speeds (rows indicated at right adjacent to heat map). Fungal DNA yields from these 60 variations in isolation methodology were calculated from Fungiquant qPCR as described in Figure 1, then scaled across all samples. Values are expressed as a heat map, with bright red signifying the highest yields and black the lowest yields across all samples.

### Effect of sample volume on community profiling

Specimen volume is thought to influence the representation of microbial complexity determined by NGS, particularly in low biomass specimens, such as urine[44], which was also suggested by our preliminary results (Figure 1C). To determine the magnitude of the effect of sample volume on microbial yields and community depth and diversity, we examined microbial profiles across a range of urine sample volumes. Large volume urine specimens from individual subjects were divided into 1, 2.5, 5, 10, 25, and 50 ml aliquots and processed in parallel according to our optimized protocol described above. Fungal yields (Figure 3A) were greatest with larger urine volumes. However, the optimal volume of initial urine was 25 ml, with 10 and 25 ml samples yielding substantially greater DNA concentrations than smaller or larger amounts. Average yields across all specimens *decreased* in 50 ml samples.

**Figure 3.**
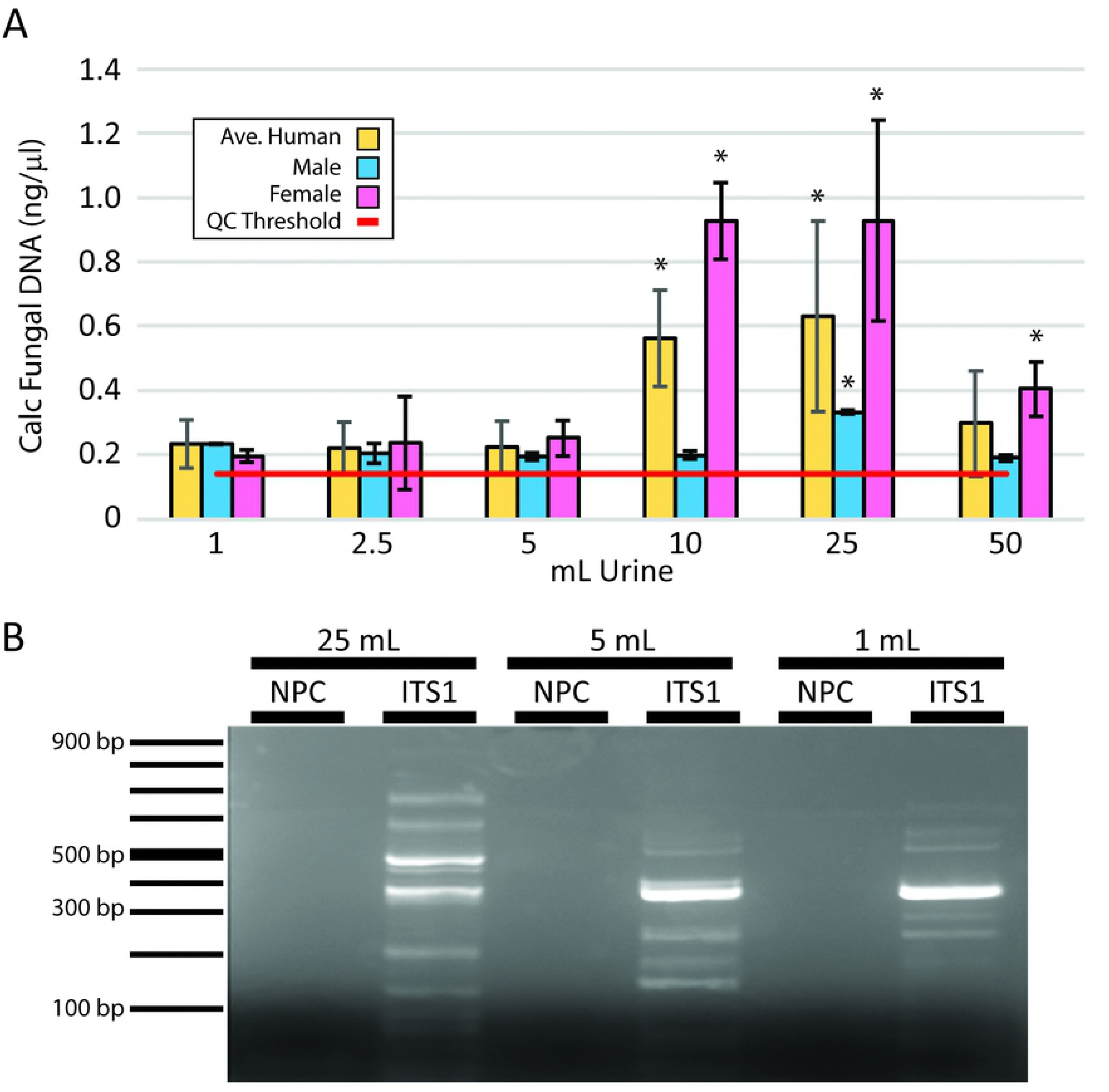
Fungal community representation is influenced by specimen volume. (A) DNA was isolated from a range of urinary volumes in male and female subjects (n=3 each) and assessed by qPCR for fungal DNA. Calculated fungal DNA concentrations were calculated by normalization to a fungal standard. The optimal concentrations were achieved using 25 ml urine specimens. *: *P*<0.05 in comparison to 1 mL yields. (B) Following fungal DNA amplification by qPCR using broad-spectrum fungal primers, products from 25, 5 and 1 ml samples were assessed by 2% agarose gel electrophoresis. Standards indicating the PCR product size are shown on the left. Each band represents unique taxa within the urinary fungal population. NPC: no primer control.

We also assessed community complexity by gel electrophoresis following PCR-based amplification of the fungal ITS1 rDNA region in which different sized products represent unique fungal taxa (Figure 3B). In comparison to sample sizes of 5 ml or less, 25 ml provided a more comprehensive representation of the range of fungal species with an increased number of bands of varying sizes representing unique taxa for larger initial sample sizes. Across all volumes, urine from male subjects consistently demonstrated lower yields. Only at the 25 ml volume were fungal DNA yields consistently above quality control thresholds.

### Effect of urine storage and centrifugation conditions on DNA extraction efficiency

In handling urine, we sporadically observed after centrifugation a substantial, sand-like pellet of varying colors. The appearance of this non-cellular pellet material was observed with refrigeration (>2 hours) of urine samples prior to processing and with high-speed centrifugation (16,000 rcf). Post-centrifugation pellets from larger urine volumes (>50 ml) also would frequently contain this material, even when pelleted at lower speeds (1500-5000 rcf) and processed at room temperature. Microscopic examination of these samples revealed a range of crystalline forms, typically amorphous urates or phosphates, depending on urinary pH. When these microcrystal salts appeared, DNA quality, as assessed by OD_260_/OD_280_ ratios, was significantly lower. Relative DNA yields were also consistently lower, suggesting that larger crystal burden interfered with DNA purification. One such post-centrifugation specimen (shown in Figure 4A) demonstrates a red-orange, sandy pellet, the “brick-layer’s dust” characteristic of amorphous urates. Confirmation of crystal composition was supported by microscopic analysis (Figure 4B) as well as chemical properties; these pellets could be dissolved by either heating to a temperature >60°C or adding sodium hydroxide. In a smaller subset of alkaline urine specimens, refrigeration or high-speed prolonged centrifugation resulted in a light-colored sandy pellet, which could be identified as amorphous phosphates by microscopy (Figure 4C). Chemical composition was confirmed by solubility in glacial acetic acid and resistance to dissolution with heating[45]. Other crystal forms were occasionally noted, such as the “envelope”-type crystals characteristic of calcium oxalate (Figure 4D inset), but these did not typically constitute any sizable portion of the crystalline material. We were able to minimize the appearance of crystalline salts through a combination of expedient processing (within 4 hours of sample acquisition), the avoidance of refrigeration, and optimization of sample size and centrifugation speed.

**Figure 4.**
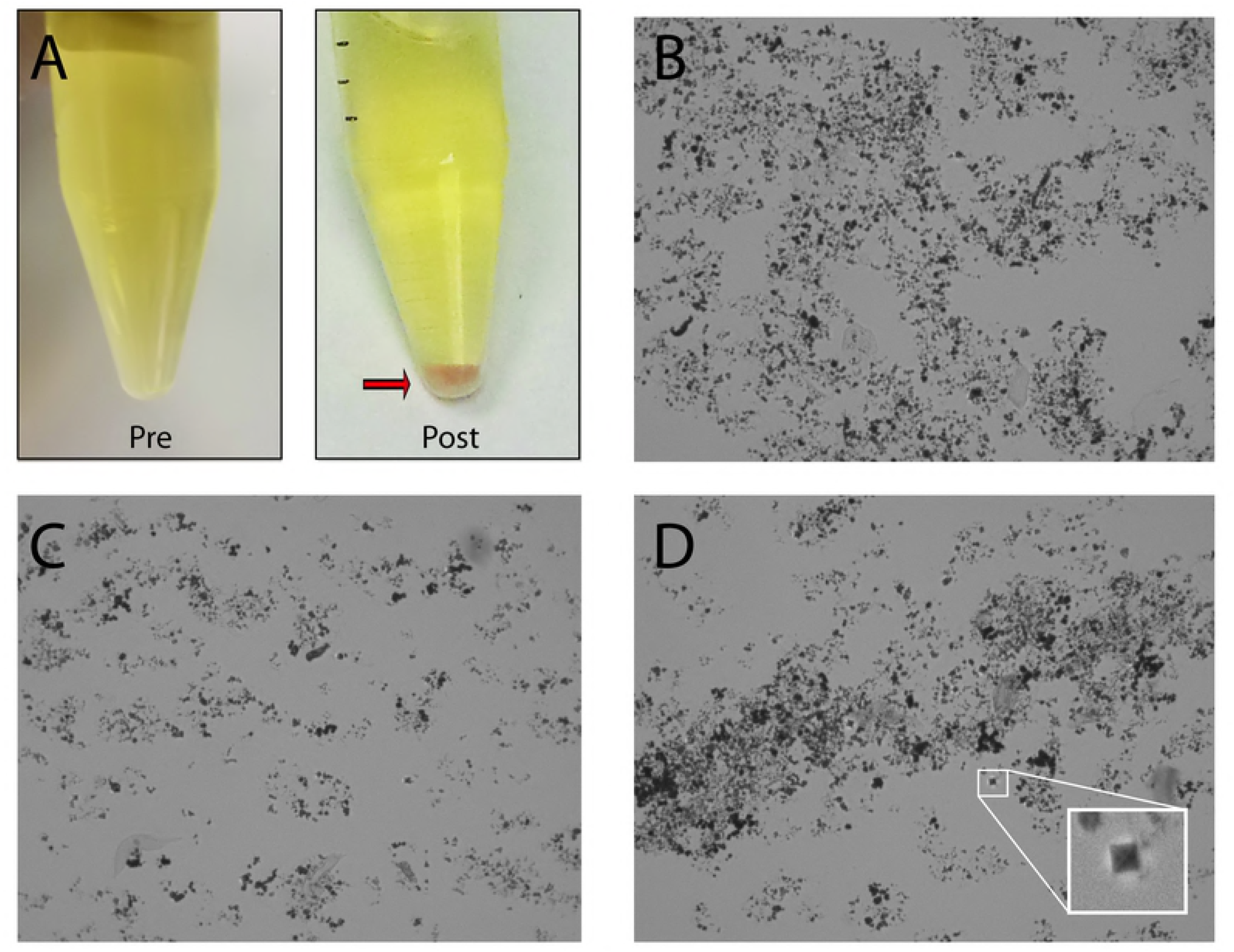
Urine storage and centrifugation conditions impact DNA extraction efficiency. In a subset of urine samples, both refrigeration and high-speed centrifugation were associated with precipitation of varying crystals that interfered with DNA purification. (A) A single urine specimen before and after refrigeration and centrifugation at 5000 rcf. In the post-centrifugation specimen, a red-orange, sandy pellet was observed after centrifugation consistent with the “brick-layer’s dust” characteristic of amorphous urates. (B) The pellet seen in A was examined by light microscopy (×400 magnification), revealing disorganized amorphous urate crystals. (C) Amorphous phosphates from alkaline urine. (D) The “envelope”-type crystals characteristic of calcium oxalate could also be identified in urine (magnified in the inset picture), but did not constitute the majority of the crystalline material.

As amorphous urates and phosphates can inhibit individual steps in DNA purification and PCR amplification, we next sought to determine if varying processing volumes could minimize any impact of these salt contaminants on DNA purification and subsequent PCR amplification. In addition, we hypothesized that smaller sample volumes might be more effectively heated for thermal disruption. After combined enzymatic treatment and mechanical cell wall disruption, we subdivided samples into varying aliquot sizes for the two boil/freeze cycles, proteinase K digestion, and DNA isolation using a DNA-binding column. Volumes ranging from 25% to 80% (100-400 μl in 50 μl increments) of the total sample lysate were applied to the spin columns before washing and DNA elution. Small sequential increases in DNA yields were seen up to 250 μl, but then plateaued, without additional increase in DNA yields with larger volumes (Supplemental Figure 1A).

**Supplemental Figure 1**. *Smaller volume sample processing and pooling increases DNA purification yields*. Standardized urine samples from 4 subjects were pelleted and processed using the optimized purification protocol for enzymatic treatment and cell wall disruption. (A) Prior to the addition of proteinase K, varying quantities of the total sample lysate were transferred to new tubes for digestion and DNA column binding. Lysate quantities ≥250 μl provided equivalent yields. (B) Prior to the addition of proteinase K, sample lysates were divided into 250 μl aliquots. Processing of a single 250 μl aliquot (No pooling) was compared to the results if the lysate was split into two aliquots and processed in parallel, then later pooled on either a single DNA-binding column and eluted as a single sample (1 column) or purified separately on two columns, eluted independently and then pooled (2 columns). Control samples were processed in parallel and did not have any input cellular material.

These data suggested that a portion of the DNA in our samples was not either effectively digested or binding to the extraction column. We therefore attempted pooling of subdivided samples; sample lysates were divided into equal halves (~250 μl) and processed in parallel before column binding. Lysates were then either pooled onto a single column in two subsequent binding steps and eluted in a single elution or bound and eluted from separate columns and pooled after elution. Pooling of two 200-250 μl aliquots on a single DNA column provided the best DNA yields (Supplemental Figure 1B).

### Our optimized method outperforms previously described and commercial DNA isolation methods

We then evaluated our method in comparison to several commercial DNA kits commonly used for microbial analysis. This optimized protocol yielded higher concentrations of DNA and greater species diversity for fungal DNA than identical samples processed with the PSP^®^ Spin Stool DNA Plus Kit, PureLink™ Microbiome DNA Purification Kit, and QIAamp DNA Stool Mini Kit (Figure 4). As the ideal goal of this method would be the simultaneous examination of both fungal and bacterial populations, we also assessed the utility of the optimized protocol in the isolation of bacterial DNA. Our protocol consistently outperformed commercial methods for the purification of fungal as well as bacterial DNA (Figure 5A). To assess the applicability of this protocol to other human commensal microbial communities, we analyzed a panel of vaginal swabs as well. Our protocol enhanced fungal and bacterial recovery from vaginal swabs significantly. While the method previously described for urine samples[5, 32] using Qiagen DNA isolation kits (Qiagen) was already better than the commercial kits tested, the optimized protocol increased the yield of fungal DNA approximately 200% (*p*<0.001) for vaginal swabs and 130% (*p*<0.005) for 30 ml urine samples. Bacterial yields differed even more profoundly, increasing yields approximately 240% for vaginal swabs and 200% for urine over levels seen with the best of previously described methods (*p*<0.001).

**Figure 5.**
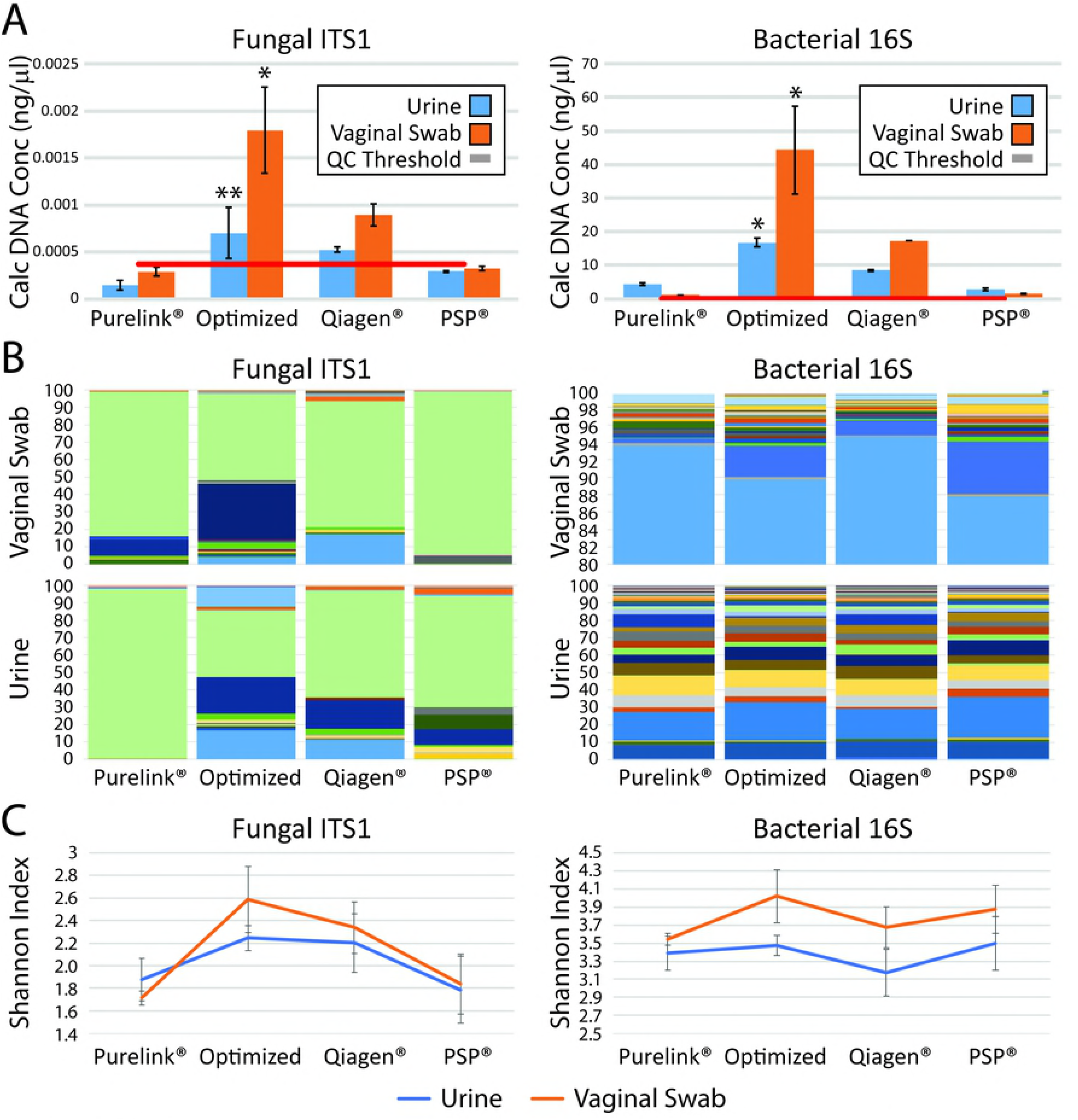
Our optimized DNA extraction method outperforms commercial methods. We compared fungal (left) and bacterial (right) extraction and characterization after our optimized protocol in comparison to three commercial DNA preparation kits using standardized urine and vaginal swab samples. (A) Individual urine specimens were divided into equal aliquots of 30 ml each. DNA was isolated from each aliquot using the specified methods; this process was repeated in quadruplicate. Samples were assessed by qPCR for fungal (left) and bacterial (right) DNA. Calculated DNA concentrations were determined by normalization to a mixed fungal and bacterial standard with a known DNA concentration. *: *P*<0.001, **: *P*<0.005. (B,C) Samples were sequenced in quadruplicate by next generation sequencing for the ITS1 (left) and 16S (right) primers. (B) The stacked bar plots represent the mean relative abundances for the fungal (left) and bacterial (right) populations in individual sequencing runs. (C) Shannon diversity indices were calculated from the microbial populations resulting from NGS for each purification method.

The improved yields translated to an improved representation of urinary microbial community diversity as assessed by NGS. Qualitatively, a community of greater richness and evenness, as measured using the Shannon Diversity Index (Figure 5C), was seen; multiple taxa were absent or underrepresented in other purification methods (Figure 5B). The optimized method consistently resulted in the highest diversity of all methods. While these differences were not statistically significant, our optimized technique provides equivalent or improved bacterial and fungal community representation across multiple biological sample types.

## Discussion

Extraction of DNA from fungal cells in urine has proven challenging for multiple reasons. Fungi are thought to be low abundance in most body sites and are structurally more robust and difficult to lyse. Multiple challenges in the identification and characterization of fungal species, such as incomplete annotation in common databases, inconsistent taxonomic classification, and variable conservation of the ribosomal locus across divisions of the fungal kingdom[46], complicate studies of fungi in any biologic niche. The combination of these problems with the technical difficulties of working with urine specimens has left previous explorations of the urinary fungal microbiota inadequate to examine anything more than a few, well-characterized species.[25] Given the experiences of others attempting fungal isolation in low biomass specimens, such as blood, we anticipated that substantial modifications of typical protocols used for the isolation of bacteria from urine would be necessary to assess adequately the fungal populations present. As optimal depth of sequencing requires the highest concentrations of DNA possible, we hypothesized that successful fungal DNA extraction for sequencing would require concentration of cellular material from larger volumes of urine, multiple disruption steps to break down fungal cell walls, and inactivation of the abundant PCR inhibitors present in urine. An iterative approach to the optimization of fungal DNA extraction confirmed these suspicions, with multiple modifications from commonly-used standard DNA extraction methods needed to provide consistent, good quality fungal DNA for sequencing-based assessments.

Sample size was very important. Approximately 40% of low volume specimens (e.g. 1 ml urine) did not provide adequate sequencing depth for analysis (<1000 reads per sample), while samples >10 ml consistently provided excellent depth of coverage in ~95% of samples. While this volume threshold had previously been suggested[44], our data provides objective confirmation that such a threshold is important for microbial analyses. Unexpected was the discovery that the best results were not associated with the largest, initial sample size, with an optimal samples size of 25-30 ml. Interestingly, centrifugation speed made a substantial difference, with slower speeds yielding better results. While both these results seem counter-intuitive, our data suggest that this decrease in fungal DNA seen with higher centrifugation speeds and larger sample volumes is due to the accumulation of amorphous crystals common in urine that interfere with DNA extraction and amplification by PCR. While it is possible that other methodologic variations, such as filter-based concentration methods or magnetic bead separations, could provide improved results, our initial attempts using these methods did not appear promising.

As anticipated, multiple cell wall disruption methods (thermal, mechanical, and enzymatic) provided much improved fungal DNA yields. An additional digestion with proteinase K was helpful at improving DNA quality as well, although the precise amount of enzyme was less important. The use of carrier DNA to enhance DNA column binding efficiency was crucial. Parallel processing of cell lysates in smaller batches with serial application of these samples to a single DNA binding column also improved yields. The individual improvements in fungal DNA extraction for each of these steps justified their inclusion into the optimized protocol.

We have also attempted multiple other variations on our protocol that are not described in this paper, such as preheating the DNA elution buffer to 37°C or performing a second elution from the DNA-binding column in a small volume, as none of these possibilities made significant differences in the resulting DNA concentrations. Our results without these additional steps were sufficient for genomic sequencing.

One drawback to these additional steps is that this protocol takes significantly more time than the available commercial kits, 150-180 min in contrast to 75-90 min. The substantial improvement in the quality and quantity of isolated microbial DNA, however, is clear, consistently providing reliable DNA for NGS analyses of microbial populations.

The samples utilized as test specimens throughout this paper were voided. Contamination from nearby sites, such as skin, urethra, and vagina (in women), can contribute heavily to the microbial content of voided samples[44]. When compared directly (Figure 3), fungal levels in samples from women were 2-3 fold higher than those seen for men. While this could reflect a difference in the urinary fungal content between genders, it may also merely reflect differences in contamination from nearby urogenital sites. As a result, this paper does not seek to make conclusions about the composition of the urinary mycobiome, but instead sought to explore the solutions needed to characterize microbial content from urine specimens. Larger scale studies, which are currently underway, using a multitude of samples will be needed to explore the urinary mycobiome. However, while the samples used in this study were voided in origin, we have since confirmed that this enhanced protocol is successful at producing sufficient quality fungal DNA to obtain good depth of sequencing from a limited number of catheterized urine samples and those obtained by suprapubic aspirate.

For microbial populations of low abundance, as presumed for the urinary tract, maximizing the quantity of template DNA for analysis is extremely important. When DNA quantities are barely in the range of detection, small variations in sample quantity or quality or even minor fluctuations in physiologic conditions may result in large misleading population shifts. If certain benign urologic conditions are associated with changes in the overall abundance of fungi in urine, as has been suggested for UCPPS,[25] then methods that fail to adequately represent the population at the lower, baseline levels will underrepresent the populations present in these circumstances. It is likely in that situation that culture-independent microbial analyses will incorrectly identify the upregulation or novel appearance of particular taxa, providing misleading conclusions about disease pathophysiology. These problems are compounded by the fact that urine composition and concentration is highly variable, even within a single individual. Certain disease conditions are associated with systematically smaller void volumes, which might also significantly bias such results. The increased DNA concentration and quality achieved using this optimized approach seek to minimize these biases and provide the most accurate results in the use of sequencing-based methods to define the urinary mycobiome.

It has been widely recognized for bacterial DNA extraction that different sample preparation and DNA extraction protocols can produce dramatically different results.[47-50] Protocols utilizing mechanical and enzymatic disruption steps have consistently given the best representations of bacterial community structure, but in no case have the obtained results provided completely accurate representations of standardized samples.[50] In fungal studies,[51] optimal conditions vary for individual fungal species; therefore, while standardized methods are generally useful for fungal and bacterial DNA extraction from biologic specimens, every method will have some bias in extraction efficiency. No single extraction method is reliable and optimal for all species in all specimens. While our results from a range of subjects and specimens confirmed the efficacy of this optimized protocol in aggregate, there were individual variations in fungal community patterns. Our optimized protocol as defined was not always the most effective for every subject assessed. The greatest variations occurred with centrifugation conditions; it is likely that for subjects for whom there is a lower urinary salt content there would be improved results with higher centrifugation speeds. Such biases are inevitable for all stages in the process of culture-independent sequencing-based identification of microorganisms. It remains important to keep these biases in mind when interpreting results, as well as to confirm results through multiple methodologies.

In conclusion, we present a method for microbial DNA isolation that results in a better representation of the overall fungal and bacterial populations, both in terms of the population diversity as well as identification of low abundance taxa that are lost with less sensitive methods. All of these benefits appear to occur without a significant loss in bacterial community representation, making this the best available method for microbial analyses of urine samples.

## Conclusion

Studies examining urinary fungal populations have been limited by the inability to consistently isolate the microbial DNA from low biomass urinary samples. This report describes an optimized protocol for the analysis of urinary fungi that is also highly effective for the concurrent analysis of urinary bacterial populations. The simultaneous and efficient extraction of fungal and bacterial DNA from urine for use in culture-independent microbial analyses is thus possible with this refined technique, providing more reliable methods for the detection and exploration of multiple microbial kingdoms from a single specimen.

## MAPP I and II Research Network

**MAPP Network Executive Committee**

**J. Quentin Clemens, MD, FACS, MSci, Network Chair, 2013-**

Philip Hanno, MD

Ziya Kirkali, MD

John W. Kusek, PhD

J. Richard Landis, PhD

M. Scott Lucia, MD

Robert M. Moldwin, MD

Chris Mullins, PhD

Michel A. Pontari, MD

**University of Colorado Denver Tissue Analysis & Technology Core**

**M. Scott Lucia, MD, Core Dir. Adrie van Bokhoven, PhD, Co-Dir**.

Andrea A. Osypuk, BS

Robert Dayton, Jr

Chelsea S. Triolo, BS

Karen R. Jonscher, PhD

Holly T. Sullivan, BS

R. Storey Wilson, MS

Zachary D. Grasmick, BS

**National Institutes of Diabetes & Digestive and Kidney Diseases**

Chris Mullins, PhD

John W. Kusek, PhD

Ziya Kirkali, MD

Tamara G. Bavendam, MD

**University of Pennsylvania Data Coordinating Core**

**J. Richard Landis, PhD, Core Dir**.

Ted Barrell, BA

Ro-Pauline Doe, BA

John T. Farrar, MD, MSCE, PhD

Melissa Fernando, MPH

Laura Gallagher, MPH, CCRP

Philip Hanno, MD

Xiaoling Hou, MS

Tamara Howard, MPH

Thomas Jemielita, MS

Natalie Kuzla, MA

Robert M. Moldwin, MD

Craig Newcomb, MS

Michel A. Pontari, MD

Nancy Robinson-Garvin, PhD

Sandra Smith, AS

Alisa Stephens-Shields, PhD

Yanli Wang, MS

Xingmei Wang, MS

**Northwestern University**

**David J. Klumpp, PhD, Co-Dir. Anthony J. Schaeffer, MD, Co-Dir**.

Apkar (Vania) Apkarian, PhD

Christina Arroyo

Michael Bass, PhD

David Cella, PhD

Melissa A. Farmer, PhD

Colleen Fitzgerald, MD

Richard Gershon, PhD

James W. Griffith, PhD

Charles J. Heckman II, PhD

Mingchen Jiang, PhD

Laurie Keefer, PhD

Robert Lloyd, PhD

Darlene S. Marko, RN, BSN, CCRC

Jean Michniewicz

Richard Miller, PhD

Todd Parrish, PhD

Frank Tu, MD, MPH

Ryan Yaggi

**University of California, LA PAIN Neuroimaging Core**

**Emeran A. Mayer, MD, Co-Dir. Larissa V. Rodríguez, MD, Co-Dir**.

Jeffry Alger, PhD

Cody P. Ashe-McNalley

Ben Ellingson, PhD

Nuwanthi Heendeniya

Lisa Kilpatrick, PhD

Cara, Kulbacki

Jason Kutch, PhD

Jennifer S. Labus, PhD

Bruce D. Naliboff, PhD

Fornessa Randal

Suzanne R. Smith, RN, NP

**University of Iowa**

**Karl J. Kreder, MD, MBA, Dir**.

Catherine S. Bradley, MD, MSCE

Mary Eno, RN, RA

Kris Greiner, BA

Yi Luo, PhD, MD

Susan K. Lutgendorf, PhD

Michael A. O’Donnell, MD

Barbara Ziegler, BA

Andrew Schrepf, PhD

Isabelle Hardy, MBA

Vince Magnotta, PhD

Brad Erickson, MD

**University of Michigan**

**Daniel J. Clauw, MD, Co-Dir.; Network Chair, 2008-2013**

**J. Quentin Clemens, MD, FACS, MSci, Co-Dir.; Network Chair, 2013-**

Suzie As-Sanie, MD

Sandra Berry, MA

Clara Grayhack,

Megan E. Halvorson, BS, CCRP

Richard Harris, PhD

Steve Harte, PhD

Eric Ichesco, BS

Ann Oldendorf, MD

Katherine A. Scott, RN, BSN

David A. Williams, PhD

**University of Washington, Seattle**

**Dedra Buchwald, MD, Dir**.

Niloofar Afari, PhD, UCSD

Tamara Bacus, BS

Todd Edwards, PhD

John Krieger, MD

Kenneth Maravilla, MD

Jane Miller, MD

Donald Patrick, PhD

Xiaoyan Qin, PhD

Stephanie Richey, BS

Rosana Risques, PhD

Kelly Robertson, BS

Susan O. Ross, RN, MN

Roberta Spiro, MS

Eric Strachan, PhD

TJ Sundsvold, MPH

Suzette Sutherland, MD

Claire C. Yang, MD

**Washington University, St. Louis**

**Gerald L. Andriole, MD, Co-Dir., PI H. Henry Lai, MD, Co-Dir., PI**

Rebecca L. Bristol, BA, BS

Robert W. Gereau IV, PhD,

Barry A. Hong, PhD, FAACP

Aleksandra P. Klim, RN, MHS, CCRC

Siobhan Sutcliffe, PhD, ScM, MHS

Joel Vetter

David G. Song

Melissa Milbrandt

Simon Haroutounian, PhD

Pooja Vijairania

Kaveri Parker (Chaturvedi)

Tran Hung

Graham Colditz, MD, PH

Vivien C. Gardner, RN, BSN

Jeffrey P Henderson, MD, PhD

Theresa M. Spitznagle, PT, DPT, WCS

Ratna Pakpahan, MHA

Aimee James PhD, MPH

Yan Yan

Marvin Epolian Langston

Barry Hong, PhD

Susan Mueller

Jan Crowley

Sherri Vogt

Scott Hultgren, PhD

Nang Nguyen, PhD

Gabriel Blasche

Chang Shen Qiu, PhD

Lori Cupps

Song Bok

**Thomas M. Hooten, MD [U. Miami]**

Lucy Grullon [U Miami]

Nadege Atis [U Miami]

**Timothy J. Ness, MD, PhD [UAB]**

Georg Deutsch, PhD [UAB]

Jan Den Hollander, PhD [UAB]

Beverly D. Corbitt, RN [UAB]

Laurence Bradley, PhD [UAB]

**Carol S. North, MD, MPE, [UTSW]**

Dana Downs, MA **[UTSW]**

**NON-RECRUITING DISCOVERY SITES**

**Cedars-Sinai Medical Center**

Jennifer Anger, MD, MPH

**Cedars-Sinai Medical Center**

Jennifer Anger, MD, MPH

James Ackerman, MA

A. Lenore Ackerman, MD, PhD

Jeena Cha, BS, CCRP

Karyn Eilber, MD

Michael Freeman, PhD

Vincent Funari, PhD

Jayoung Kim, PhD

Jennifer Van Eyk, PhD

Wei Yang, PhD

**Queens University**

**Harvard Medical School/**

Boston Children’s Hospital

**Masha A. Moses, PhD, Dir**.

Andrew C. Briscoe,

David Briscoe, MD

Adam Curatolo, BA

John Froehlich, PhD

Richard S. Lee, MD

Monisha Sachdev, BS

Keith R. Solomon, PhD

Hanno Steen, PhD

**J. Curtis Nickel, MD, FRCSC, Dir**.

**Stanford University**

**Sean Mackey, MD, PhD, Dir**.

Epifanio Bagarinao, PhD

Lauren C. Foster, BA

Emily Hubbard, BA

Kevin A. Johnson, PhD, RN

Katherine T. Martucci, PhD

Rebecca L. McCue, BA

Rachel R. Moericke, MA

Aneesha Nilakantan, BA

Noorulain Noor, BS

Garth D. Ehrlich, PhD, [Drexel COM]

